# Early Flowering 3 (ELF3) inhibits hypocotyl phototropism in light-grown Arabidopsis seedlings

**DOI:** 10.1101/2025.04.09.647958

**Authors:** Geoffrey Cobb, Johanna Krahmer, Ganesh M. Nawkar, Alessandra Boccaccini, Christian Fankhauser

## Abstract

Phototropic bending of plants towards a light source allows them to position their photosynthetic tissues to optimize light capture. In light-grown (deetiolated) *Arabidopsis* seedlings, phototropic bending of the hypocotyl is inhibited by light with a high red:far-red ratio (HRFR) and high levels of blue light (HBL). This occurs via activation of the phytochrome B (phyB) and cryptochrome 1 (cry1) photoreceptor signaling pathways. Both phyB and cry1 act upstream of PHYTOCHROME INTERACTING FACTOR (PIF) transcription factors, which are required for hypocotyl bending in light-grown seedlings. Presently, it is not known whether other pathways are involved in the inhibition of PIF-mediated phototropism in light-grown seedlings. To address this, we conducted a screen to identify mutants with increased phototropic bending relative to wild type in HRFR+HBL conditions. Through this screen, we identified EARLY FLOWERING 3 (ELF3), a member of the Evening Complex (EC), as a key inhibitor of phototropic bending in green seedlings. We show that both ELF3 and LUX, another component of the EC, inhibit phototropic bending upstream of PIF4/PIF5. Furthermore, we show that phototropic bending in *Arabidopsis* seedlings is subject to circadian regulation in an ELF3-dependent manner. Finally, we provide evidence that ELF3 in the grass *Brachypodium distachyon* also affects phototropism but in an opposite way than in *Arabidopsis*.

**One sentence summary:** ELF3 inhibits hypocotyl phototropism in de-etiolated seedlings growing in favorable light conditions.

## Introduction

Plants in the wild must often compete with neighboring plants for resources such as water, nutrients, and light. In canopy shade conditions, where a plant is completely overtopped by its neighbors, the ratio of red to far-red light is lower (LRFR conditions), and the level of blue light is also lower (LB conditions) than in full sunlight (Fiorucci and Fankhauser, 2017). In these environments, plants may bend towards available sources of light in order to maximize their exposure. In light-grown (deetiolated) *Arabidopsis* seedlings, this phototropic bending requires the PHYTOCHROME INTERACTING FACTORS (PIFs) 4, 5, and 7 (Goyal et al., 2016; Boccaccini et al., 2020). PIF4/5/7 are transcription factors that promote the expression of *YUCCA* auxin biosynthesis genes, which are themselves required for bending (Goyal et al., 2016).

In sunny conditions, hypocotyl phototropism is weak in de-etiolated Arabidopsis seedlings, presumably due to sufficient access to light (Iino, 2001; Boccaccini et al., 2020). In this case, the high red:far-red ratio (HRFR) and higher levels of blue light (HB) activate phyB and cry1 photoreceptor signaling pathways, which results in the repression of the PIFs, thereby inhibiting phototropism (Goyal et al., 2016; Boccaccini et al., 2020). PIF4/5/7 thus form a critical link between sensing of a plant’s light environment and growth responses such as phototropism. Whether PIFs are regulated in additional ways to control their function in hypocotyl phototropism remains unknown.

The Evening Complex (EC) is a protein complex that plays a critical role in the function of the circadian clock in plants by repressing the expression of key genes around the end of the light photoperiod (Huang and Nusinow, 2016). The evening complex in *Arabidopsis* has three components: EARLY FLOWERING 3 (ELF3), EARLY FLOWERING 4 (ELF4), and LUX ARRYTHMO (LUX) (Huang and Nusinow, 2016). While the evening complex has not previously been demonstrated to affect phototropic bending, it is known to repress the transcription of PIFs at the end of long days, resulting in a daily cycle of PIF transcript levels (Nusinow et al., 2011; Murcia et al., 2022). Protein levels of PIF4/5/7 follow their own transcript levels closely (Galvao et al., 2019; Murcia et al., 2022), indicating that the evening complex likely plays a key role in maintaining a daily cycling of PIF protein levels. Additionally, ELF3 is known to repress the activities of PIF4 and PIF7 independently of the evening complex via direct protein-protein interaction, which prevents PIFs from binding to their target genes (Nieto et al., 2015; Jiang et al., 2019). As important regulators of PIFs, the evening complex members are good candidates for possible regulators of phototropism in light-grown seedlings.

Historically, mutant screens in dark-grown (etiolated) *Arabidopsis* seedlings have successfully identified many components necessary for phototropic bending. For example, the blue light photoreceptor phot1 was identified in several screens for mutants with reduced capacity for hypocotyl bending in response to unilateral blue light (Khurana and Poff, 1989; Liscum and Briggs, 1995). NPH3 and RPT2, important regulators of phototropic bending and direct interactors of PHOT1, were also identified in such screens (Okada and Shimura, 1992; Liscum and Briggs, 1995), as were the cryptochromes CRY1 and CRY2 (Ahmad et al., 1998). Phot1, NPH3, and cry1 have since been demonstrated to be important regulators of phototropism in light-grown seedlings as well as dark-grown seedlings (Goyal et al., 2016; Boccaccini et al., 2020). However, phototropic bending in dark-grown seedlings does not require PIF4/5/7, meaning that these screens are insufficient to identify regulators of PIF-mediated phototropism in light-grown seedlings (Goyal et al., 2016).

To address this, we initiated a mutant screen to identify negative regulators of phototropism in light-grown *Arabidopsis* seedlings. Through this screen, we identified mutants defective in phytochrome and cryptochrome signaling, confirming the importance of these pathways to the control of phototropism. We also identified a nonsense allele in *ELF3* which conferred increased phototropic bending over wild-type (WT) in our screen conditions. We demonstrate that ELF3 and LUX act upstream of PIF4/5 to inhibit bending. Furthermore, we show that bending is subject to diurnal regulation whose rhythm rapidly attenuates upon transfer to constant light conditions. Finally, given that ELF3 and the PIFs are very highly conserved among land plants, we performed an initial characterization of the role of the *ELF3* gene in regulating phototropism in the grass *Brachypodium distachyon*.

## Results

### A mutant screen for negative regulators of phototropism in light-grown Arabidopsis seedlings

To identify negative regulators of phototropic bending in light-grown seedlings, we first determined growth conditions under which wild type (Col-0) seedlings do not bend towards a lateral light source, while a loss-of-function mutant in a known inhibitor of phototropic bending (*phyB-9*) is able to bend. The precise conditions used for the screen are detailed in the Materials and Methods.

Using these growth conditions, we screened an M2 population of EMS-mutagenized seeds for mutants with increased phototropic bending relative to wild type. In total, we tested approximately 27,000 individual M2 seedlings, sourced from 126 seed pools, with each seed pool containing seeds from 35 M1 plants. A small number of seedlings with increased phototropic bending were grown for at least 1 additional generation and retested, then backcrossed to WT. An overview of the testing regime is shown in Figure S1B. Bending phenotypes were then assessed in the F1 and F2 backcross to determine their segregation pattern.

Using this method, we identified candidate lines with reproducible bending phenotypes, several of which are summarized in Table 1. Because phyB and cry1 are already known to negatively regulate bending (Goyal et al., 2016; Boccaccini et al., 2020), we performed a series of tests to determine whether any of our mutant lines were impacted in the phytochrome or cryptochrome signaling pathways. *phyA, phyB*, and *cry1* loss-of-function mutants have long hypocotyls in monochromatic far-red light, monochromatic red light, and monochromatic blue light, respectively (Neff and Chory, 1998). With this in mind, we assessed the hypocotyl lengths of our candidates in monochromatic red, far-red, and blue light. All of the candidate lines tested were determined to have long hypocotyls in at least one of these light conditions (Summarized in Table 1).

**Table 1.** Summary of candidates identified in the mutant screen.

| Line ID number | Hypocotyl length phenotype in monochromatic light (M3 seedlings) | Western blot result (M4 seedlings) – See Figure S5. | Sequencing result | Possible explanation for bending phenotype |
| --- | --- | --- | --- | --- |
| I040123 | long in R + FR | PHYA expressed in light and dark |  | Possible phytochrome chromophore mutant |
| C091602 | long in R + FR | PHYA expressed in light and dark |  | Possible phytochrome chromophore mutant |
| B121320 | long in B | CRY1 is not expressed |  | CRY1 is not expressed |
| C120501 | long in B | CRY1 is not expressed |  | CRY1 is not expressed |
| C061423 | Long in B | Expresses CRY1 | Substitution: R525K in *CRY1*. | Probable loss-of-function *cry1* mutant (Ahmad et al., 1995) |
| C041719 | Long in R | Expresses PHYB | Substitution: G1039R in *PHYB*. | Probable loss-of-function *phyB* mutant (Krall and Reed, 2000) |
| B111018 | Long in R | Expresses CRY1, PHYA, and PHYB | Premature stop codon at position 565 in *ELF3*. | Probable loss-of-function *elf3* mutant (Hicks et al., 2001) |

If a candidate was found to have a hypocotyl phenotype which mimicked a known photoreceptor mutant, we conducted western blots to determine whether that photoreceptor was expressed. In this way, we determined that two of our candidates (B121320 and C120105) which have long hypocotyls in blue light do not express CRY1 protein (Table 1, Figure S5).

One of our candidates (C061423) had long hypocotyls in blue light but did express CRY1 protein (Table 1, Figure S5). Sequencing of the CRY1 locus in this line revealed a substitution R525K in the C-terminal region of the protein. Interestingly, substitutions in this region are already known to result in a loss of CRY1 function (Ahmad et al., 1995), making this a likely explanation for the increased bending phenotype of this candidate. In a similar way, we identified a candidate (C041719) with long hypocotyls in red light, which expressed PHYB protein (Table 1, Figure S5). Sequencing of the phyB locus in this line revealed a substitution G1039R in the C-terminal histidine kinase domain. Mutations in this region are known to result in loss of phyB function (Krall and Reed, 2000).

Some candidates (I040123 and C091602) had long hypocotyls in both red and far-red light, indicating possible disruption of both phyB and phyA functions simultaneously. Insensitivity to both red and far-red light is a characteristic of mutants which are impaired in phytochrome chromophore biosynthesis (Goto et al., 1993). In such mutants, phyA is stable when dark-grown seedlings are exposed to light (Parks et al., 1989; Parks and Quail, 1991). Therefore, we conducted a western blot to compare phyA levels in dark-grown and white-light-exposed seedlings in our candidate lines. We found that these candidates possess photostable phyA (Table 1, Figure S5).

### A nonsense allele in *ELF3* confers increased phototropic bending

One of the candidates that we identified in our screen, B111018, exhibited increased phototropic bending over wild type in our screen conditions (Figure 1A), long hypocotyls in monochromatic red light (Figure 1B), and slightly longer hypocotyls in monochromatic far-red and blue light (Figure S2). Candidate B111018 still expressed PHYB protein and possessed photolabile PHYA (Table 1, Figure S5). To further characterize candidate B111018, we tested it for altered leaf elevation (hyponasty), which is a process that depends on PIF4/5/7 and which follows a circadian pattern (Michaud et al., 2017). Interestingly, B111018 exhibited almost completely arrhythmic leaf hyponasty in free-running light conditions, mimicking an *elf3-1* control (Figure 1C) (Hicks et al., 1996). Therefore, we sequenced the *ELF3* locus in B111018. This revealed the presence of a premature stop codon at position 565 (Figure 1D). Encouragingly, there are several characterized nonsense mutations in the ELF3 gene, which all result in loss-of-function phenotypes (Hicks et al., 2001). Our mutant is notable because the stop codon occurs relatively close to the C-terminus. For reference, the next-closest nonsense mutation to the C-terminus (*elf3-1*) has a stop codon at position 341 (Hicks et al., 2001).

**Figure 1.**
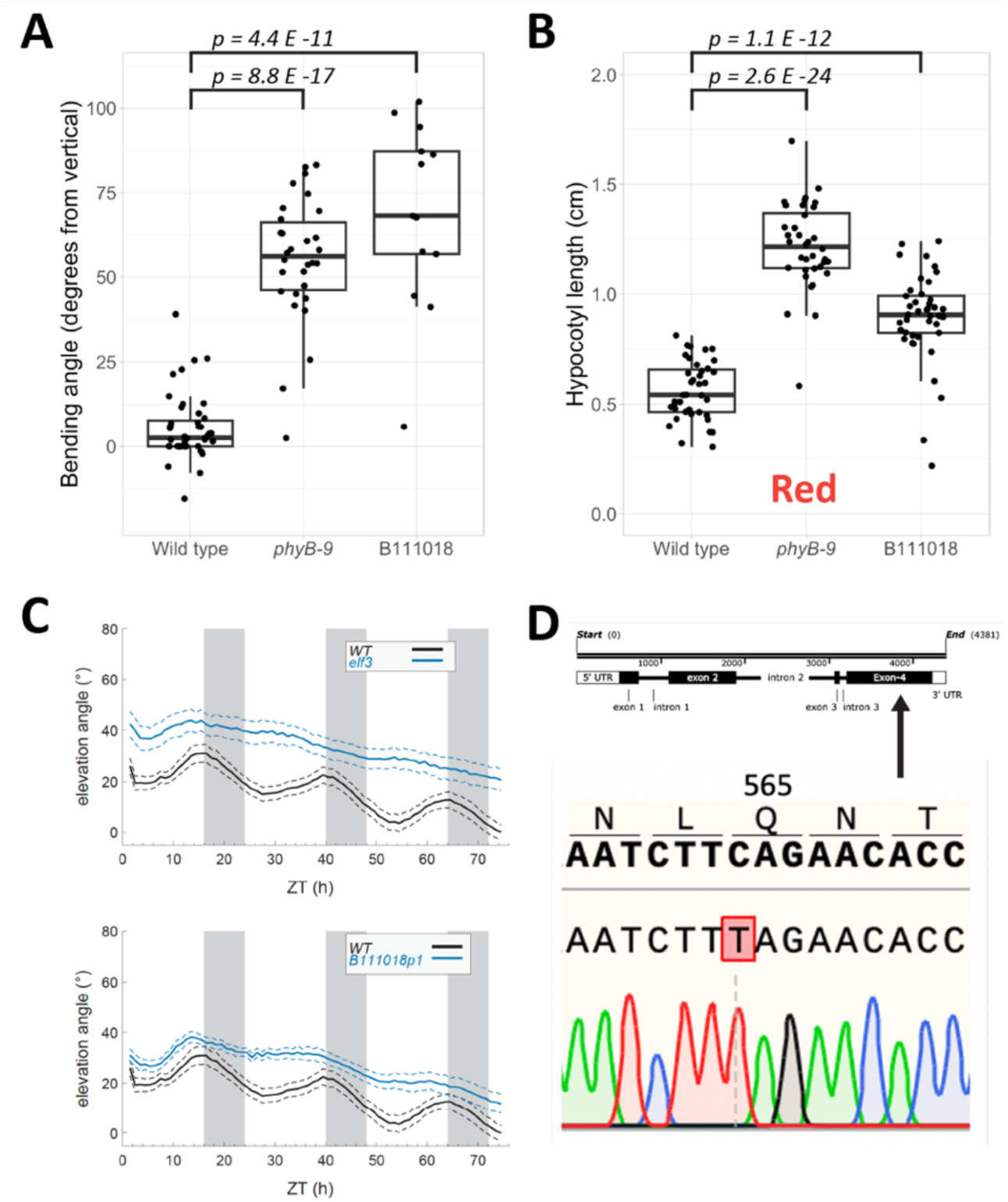
Initial characterization of the B111018 phototropic bending mutant. **(A)** Quantification of phototropic bending in B111018 seedlings grown in our screen conditions. Points represent the bending angles of individual seedlings; boxplots show the median bending angle and quartiles. For this experiment, seeds of candidate B111018 were from the M3 generation **(B)** Hypocotyl lengths of 4-day old seedlings grown in constant monochromatic red light at 14 µmol m^2^ s^−1^. Seeds from B111018 were from the M3 generation. Note: p-values in **(A)** and **(B)** are derived from Welch’s t-test. **(C)** Leaf elevation angle (angle from leaf base to tip, relative to horizontal) in free-running light conditions of the first two true leaves on long-day entrained WT, *elf3-1,* and B111018 rosettes. B111018 plants were from the M4 generation. Plants were entrained in long days (16L/8D) and were 2 weeks old upon transfer to constant light for observation. In the graphs, the solid lines represent mean leaf elevations over time, and the dotted lines indicate the 95% confidence interval around the mean estimates. Plants were measured about once per hour. In the top part of panel **(C)**, data from 13 wild-type and 13 *elf3* plants are shown. In the bottom part of the panel, data from the same wild-type individuals is shown alongside data from 12 individuals of B111018. **(D)** Schematic showing the location of the nonsense mutation in the *ELF3* locus in candidate B111018. This panel was created using Snapgene.

Next, we backcrossed candidate B111018 to wild type and observed phototropic bending in F1 and F2 seedlings. Bending in the F1 seedlings was comparable to WT, indicating that the mutation responsible for the increased bending is recessive (Figure S3A). Roughly one quarter (17/69) of the F2 seedlings bent more than the strongest-bending seedling in the wild-type control, indicating mendelian segregation of the increased bending trait (Figure S3B).

A dCAPS genotyping strategy (Neff et al., 1998) was developed to differentiate between wild-type plants and plants carrying the B111018 *elf3* mutation. This genotyping strategy was used to confirm that four F1 plants from our backcross were heterozygous for the B111018 *elf3* mutation (Figure S3C). We also genotyped four F2 seedlings which exhibited the increased bending phenotype and compared them to four F2 seedlings which lacked this phenotype. The selected F2 seedlings with increased bending were found to be homozygous for the B111018 *elf3* mutation, while those without the phenotype were found to be either homozygous for the wild-type allele or heterozygous (Figure S3D). Hence, the B111018 *elf3* mutation appears to be associated with the increased bending phenotype.

### ELF3 and LUX act upstream of PIF4/5 to control phototropic bending

ELF3 is a component of the evening complex, a transcriptional regulator which also includes ELF4 and LUX (Huang and Nusinow, 2016). Therefore, we assessed phototropic bending in the *elf3-2, elf4-101,* and *lux-4* mutants in our screen conditions. Consistent with the outcome of our screen, the *elf3-2* allele exhibited increased phototropic bending over wild type (Figure 2). Furthermore, the phototropic bending in the *elf3 pif4 pif5* triple mutant resembled that of wild-type seedlings, suggesting that ELF3 acts upstream of PIF4/PIF5 to control bending. The *lux* mutant showed similar behavior, with increased phototropic bending relative to wild type that was suppressed in both the *pif4* and *pif5* single mutant backgrounds (Figure 2). Surprisingly, while the bending in the *elf4* mutant appears to be slightly higher than wild type (Figure 2), this result was not always reproducible.

**Figure 2.**
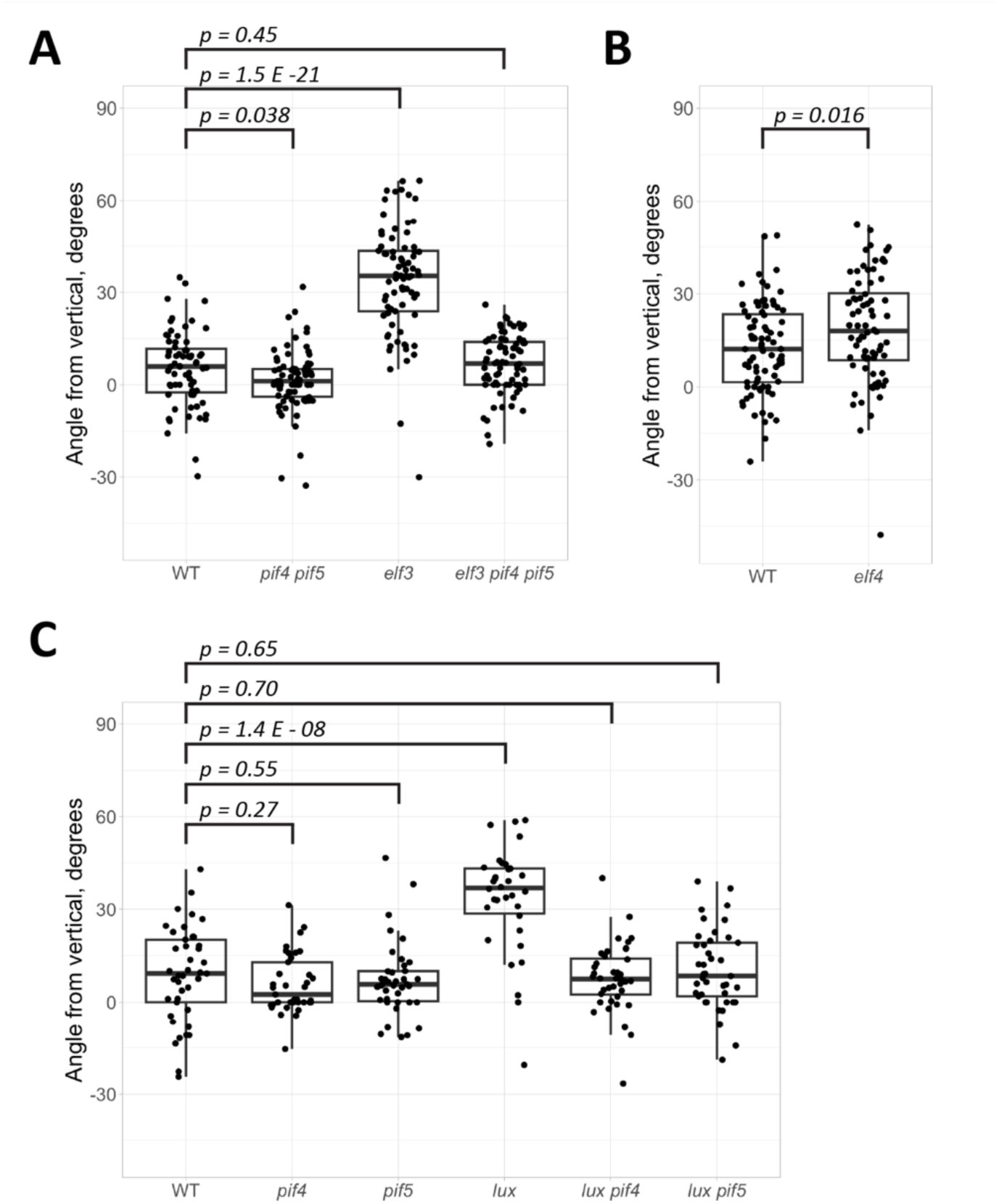
Bending assay results with *Arabidopsis* evening complex mutants. Points represent the bending angles of individual seedlings; boxplots show the median bending angle and quartiles. The p-values shown are derived from Welch’s t-test. **(A)** Bending in the *elf3, pif4 pif5,* and *elf3 pif4 pif5* mutants. All lines in this panel are in the pTOC1::LUC background. **(B)** Bending in the *elf4* mutant. **(C)** Bending in various *lux* and *pif4/pif5* mutant combinations. All lines in this panel are in the CAB2::LUC background (Nusinow et al., 2011).

### Phototropic bending potential in *Arabidopsis* seedlings follows a circadian rhythm

The evening complex is essential to the function of the circadian clock in *Arabidopsis* (Huang and Nusinow, 2016). The realization that components of the evening complex regulate phototropism indicates that bending may be subject to circadian control. Indeed, a previous study demonstrated that long day-entrained potato plantlets transferred to constant light maintain some rhythmicity in their ability to bend at different times of day, though this rhythm attenuates as the duration of exposure to constant light increases (Vinterhalter et al., 2015).

With this in mind, we wanted to test whether phototropic bending was also subject to circadian control in *Arabidopsis* seedlings. To observe strong bending in light-grown wild-type seedlings, it is necessary to grow them in combined low red:far-red (LRFR) and low blue (LB) light. Therefore, we grew seedlings in this condition under long days, at 21°C, for 4 days. At dawn on the 5^th^ day, seedlings were transferred to constant LRFR+LB light. At regular 4-hour intervals over a time course of 48 hours, subsets of seedlings were moved to a new incubator and exposed to a lateral blue light stimulus using a blue LED array. The seedlings were then photographed after 4 hours of lateral blue light exposure. A schematic of the light treatments given to each batch of seedlings is shown in Figure 3A.

**Figure 3.**
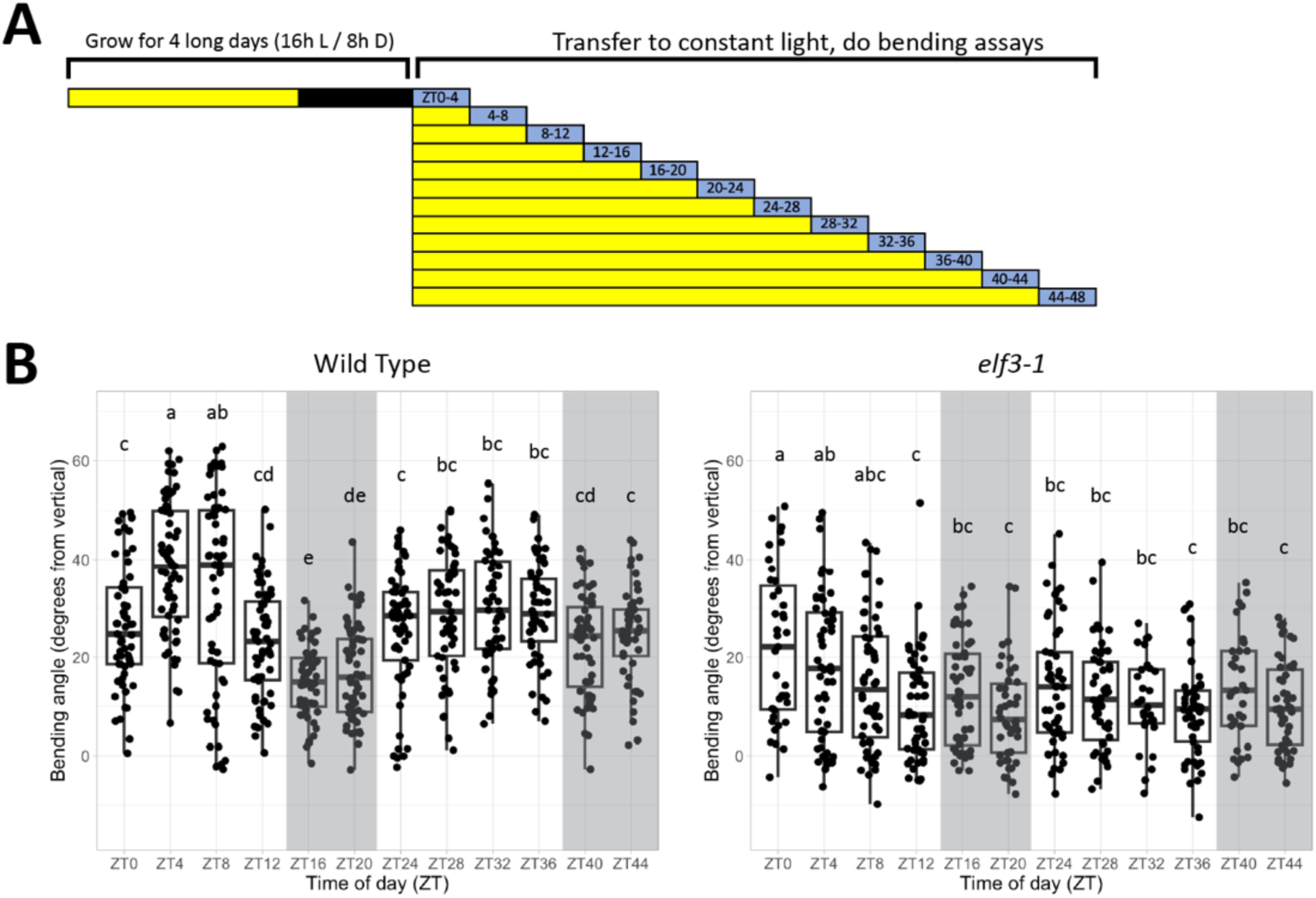
Phototropic bending at different times of day in long-day-entrained seedlings. **(A)** Schematic showing the light treatments over the course of the experiment. Seedlings were grown for 4 days in long days (16 h light / 8 h dark) at 21°C in combined LRFR+LB light. Bending assays were then conducted every 4 hours over a 48-hour timecourse, starting at dawn on the 5^th^ day. In this schematic, yellow represents combined LRFR+LB light, black represents night, and blue represents combined LRFR+LB light supplemented with lateral blue light. **(B)** Hypocotyl bending of long day-entrained seedlings transferred to constant light conditions and given lateral blue light following different durations of constant light exposure. Points represent the bending angles of individual seedlings; boxplots show the median bending angle and quartiles. Letter groupings are assigned from the results of Tukey’s HSD test with a significance level of 0.05.

We observed a peak in the bending potential of wild-type seedlings around the middle of the first subjective day (Figure 3B). Bending decreased upon onset of the first subjective night, but subsequently increased in anticipation of the second subjective day (Figure 3B). It then stabilized through the evening and into the second subjective night. By contrast, no such pattern was observed in *elf3* mutant seedlings (Figure 3B). Instead, bending in *elf3* seedlings decreased relative to the first few timepoints, then stabilized. These data confirm the involvement of the circadian clock in the emergence of rhythmic bending potentials.

### Phototropic bending in an *elf3* mutant of *Brachypodium distachyon*

Orthologs of the *PIFs* and of *ELF3* are highly conserved amongst land plants (Huang et al., 2017; Possart et al., 2017; Jiang et al., 2022). Recently, efforts have been made to characterize the functions of both the *PIF* and *ELF3* genes in the model plant *Brachypodium distachyon,* a monocot (Bouche et al., 2021; Hoang et al., 2021; Jiang et al., 2022; Gao et al., 2023). With this in mind, we wanted to determine whether ELF3 inhibits phototropism in light-grown *Brachypodium* seedlings, as it does in *Arabidopsis.* To test this, we grew wild-type and *elf3* mutant *Brachypodium* seedlings on 1/2 MS medium in the dark for 4 days, then moved them to long days in white light. On the second day in the light, the plants were put into black boxes which only allowed light to enter from one side. Bending of the seedlings towards the lateral light (defined as the angle of the coleoptile or tip of the first leaf relative to vertical) was monitored over several successive days.

Interestingly, we observed a tendency of the *elf3* mutant to bend less than the WT, which is the opposite of the bending phenotype for the *elf3* mutant in light-grown *Arabidopsis* seedlings (Figure 4A). Though bending was weak overall and quite variable for both genotypes, this difference was reproducible. To determine whether the *Brachypodium elf3* mutant has a general bending defect which could account for this, we also compared the bending phenotypes of dark-grown wild-type and *elf3* seedlings exposed to 2 µmol m^−2^ s^−1^ lateral blue light. In these conditions, bending between the wild type and *elf3* mutant was comparable, suggesting that the reduced bending phenotype of *elf3* is specific to light-grown seedlings (Figure 4B).

**Figure 4.**
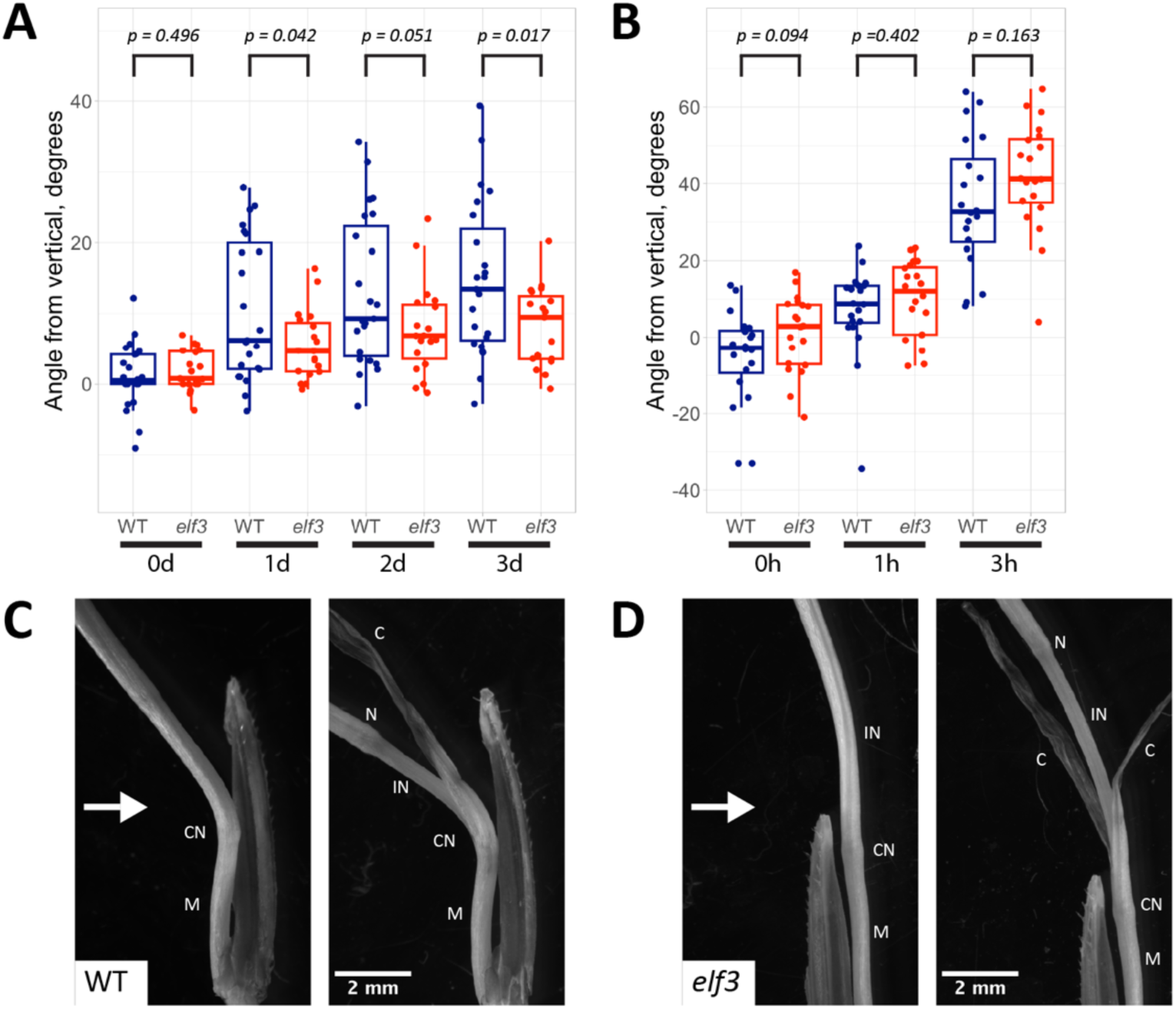
Phototropic bending in *Brachypodium* plantlets. **(A)** Phototropic bending in light-grown *Brachypodium* seedlings over the course of 3 days. Points represent the bending angles of individual plantlets; boxplots show the median bending angle and quartiles. P-values are derived from Welch’s t-test. **(B)** Phototropic bending in dark-grown *Brachypodium* seedlings exposed to 2 µmol m^−2^ s^−1^ lateral blue light. P-values are derived from Welch’s t-test. **C)** Photograph of a wild-type *Brachypodium* seedling, showing the tissues involved in phototropic bending. The direction of the lateral light source is denoted by the white arrow. The left panel shows an intact seedling, while the right panel shows the same seedling with the coleoptile peeled back to reveal underlying structures. The labels denote the following structures: mesocotyl (“M”), coleoptile node (“CN”), first internode (“IN”), second node (“N”), and the coleoptile (“C”). **(D)** Photograph of an *elf3* mutant *Brachypodium* seedling, intact (left) or with the coleoptile peeled back (right).

Because the anatomy of dicots (e.g. *Arabidopsis*) and monocots (e.g. *Brachypodium*) is markedly different, we conducted light microscopy to determine which tissues were responsible for the phototropic bending in light-grown *Brachypodium* plantlets. When seedlings are young, it is difficult to distinguish between bending of the coleoptile and other seedling structures, because they are enveloped by the coleoptile. Therefore, we observed plantlets that had been exposed to lateral light for an extended period (8 days). At this stage, the plantlets generally have a well-developed stem and 2 visible leaves. In these specimens, much of the bending seems to occur at either the coleoptile node or the first node above it (See example in Figure 4C). Curvature of the first internode region may also contribute to the bending, though this is not always present (See example in Figure 4D). This suggests that phototropism in light-grown *Brachypodium* is a complex behavior that can involve growth responses across multiple organs.

## Discussion

Positive phototropism in plants is thought to be important for allowing them to optimize their light exposure in light-limited environments (Iino, 2001; Fiorucci and Fankhauser, 2017). In light-grown *Arabidopsis* seedlings, phototropism of the hypocotyl is inhibited by the phytochrome and cryptochrome signaling pathways, which repress the action of PIFs 4/5/7, thus preventing bending in conditions when it is not needed (Goyal et al., 2016; Boccaccini et al., 2020). In this study, we sought to identify additional negative regulators of phototropism in light-grown seedlings using a forward genetics approach. We identified several candidate lines with increased phototropic bending relative to wild type in our screen conditions. Most of these lines were subsequently found to be defective in various aspects of phytochrome or cryptochrome signaling (Table 1), thus highlighting the critical importance of these signaling pathways for the regulation of phototropism. One of our candidates possesses a nonsense mutation at the *ELF3* locus. Because ELF3 is also known to act antagonistically to the PIFs, the discovery of this mutant further emphasizes the central role that the PIFs play in regulating phototropism in light-grown seedlings.

ELF3 is a member of the tripartite evening complex (EC), which is known to be important for the diurnal rhythm of *PIF4/5* transcription (Nusinow et al., 2011; Murcia et al., 2022). This complex contains two other members, ELF4 and LUX. LUX is a transcription factor and is essential for allowing the EC to interact with its target promoters (Helfer et al., 2011; Silva et al., 2020). While the molecular function of ELF4 is somewhat less clear, it has been proposed to enhance the DNA binding affinity of the EC (Silva et al., 2020), and is also required for normal EC function (Nusinow et al., 2011). With this in mind, we decided to test the bending phenotypes of the *lux* and *elf4* mutants. The *lux* mutant exhibited increased bending over wild type in our screen conditions, mimicking the phenotype of the *elf3* mutant (Figure 2). Our data is therefore consistent with the EC modulating hypocotyl phototropism in light-grown seedlings. Moreover, given the known function of the EC in regulation of PIFs, we show that the enhanced phototropic response of *elf3* and *lux* depends on the PIFs. In contrast to *elf3* and *lux*, the *elf4* mutant did not reproducibly bend more than wild type (Figure 2). The *elf4-101* allele used in our study was previously reported to have undetectable transcript levels, indicating that it is likely to be a true null (Khanna et al., 2003). As such, residual *ELF4* expression probably does not account for this observation. Therefore, we propose two alternative explanations. First, *Arabidopsis* possesses several homologs of *ELF4* known as *ELF4-LIKE* (*EFL*) genes, which are reported to act redundantly to control flowering time (Lin et al., 2019). We hypothesize that these genes might also act redundantly to regulate phototropism via the EC. Second, we note that the *elf4* mutant are reported to have stronger phenotypes in short days than in long days (Doyle et al., 2002; Hazen et al., 2005). Our bending assays were conducted with seedlings grown in long days. Therefore, it is possible that a phenotype for *elf4* mutants might be detectable in short days. In summary, our data indicate the EC limits hypocotyl phototropism in light-grown seedlings by modulating the PIFs.

The role of ELF3 and LUX in modulating the strength of phototropic bending suggest that phototropism is regulated in a circadian manner. We tested this in shade-mimicking conditions when hypocotyl phototropism is robust. Seedlings grown in shaded long days and subsequently transferred to constant light conditions maintain a rhythmic bending potential, with the maximum bending response occurring in the middle of the subjective day (4-8h after subjective dawn) and decreasing during the subjective night (Figure 3B). Notably, this pattern also correlates with published data about PIF4/5/7 protein levels, which themselves cycle in a circadian manner (Nozue et al., 2007; Galvao et al., 2019). Consistent with circadian-regulated phototropism potential, this response was lost in the *elf3* mutant. Lack of circadian rhythmicity in *elf3* mutants is also observed for other phenotypes including: leaf (Figure 1C) and cotyledon movements, and hypocotyl growth (Nozue et al., 2007). Importantly, *elf3* mutants are also known to exhibit altered circadian patterns of *PIF* transcription and PIF protein levels, further suggesting that cycling of PIF protein levels might be the mechanism responsible for the circadian gating of bending potential (Nusinow et al., 2011; Murcia et al., 2022; Zhu et al., 2022).

As previously discussed, the rhythmic bending potential in wild-type *Arabidopsis* seedlings closely resembles that of potato plantlets (Vinterhalter et al., 2015), indicating that this behavior may be found across diverse plant lineages. Interestingly, the amplitude of this rhythm decreases after the first subjective day of free-running light conditions in both *Arabidopsis* and potato (Figure 3B) (Vinterhalter et al., 2015). This suggests that constant entrainment by day/night cycles is required to maintain the rhythmic bending potential in the long term. In our experiments, the attenuation of this pattern on the second subjective day in constant light is likely exacerbated by the fact that our seedlings were grown in LRFR conditions, which is known to decrease the amplitude of circadian rhythms in free-running conditions (Fraser et al., 2021). However, we were still able to see evidence for circadian regulation of phototropism despite this limitation.

*ELF3* homologs are conserved between the dicots (including the model species *Arabidopsis thaliana*), and the monocots (including the model species *Brachypodium distachyon*), which together make up the two largest groups within the flowering plants (Huang et al., 2017). Furthermore, complementation studies suggest that the *ELF3* homolog in *Brachypodium* functions similarly to the native gene in *Arabidopsis* (Huang et al., 2017), and recent research indicates that the evening complex functions similarly in *Brachypodium* as it does in *Arabidopsis* in the control of flowering time (Gao et al., 2023). Therefore, we hypothesized that ELF3 might also regulate phototropism in monocots. To test this, we conducted bending assays with a previously described *Brachypodium elf3* mutant (Bouche et al., 2021). Bending in the *Brachypodium elf3* mutant was actually slightly lower than that of WT, and overall bending was relatively weak in both lines (Figure 4A). The reasons for this difference in phototropic behavior between *Arabidopsis* and *Brachypodium elf3* mutants are not clear. We found that phototropic bending in *Brachypodium* plantlets is derived largely from the nodes (Figure 4C), which are not analogous structures to the *Arabidopsis* hypocotyl. As such, the organs and tissue types involved in the bending response differ between both species. Notably, gravitropic bending (bending in response to gravity) has already been shown to require differential growth within the nodes of several other monocot species (Zhang et al., 2011; Clore, 2013), indicating that node-based movements may be a common feature of tropic behaviors in monocots. Moreover, there is some evidence to suggest that the roles of the PIFs may differ between *Arabidopsis* and *Brachypodium.* RNAi knockdown of *BdPIL1* and *BdPIL3,* which are orthologs of *Arabidopsis PIF1* and *PIF3*, results in longer stem internode lengths and lower leaf chlorophyll content in *Brachypodium* (Hoang et al., 2021). By contrast, analysis of various *pif* mutants in *Arabidopsis* suggests that they have the opposite effect on both phenotypes. For example, the *pif1/3/4/5* quadruple mutant in *Arabidopsis* has reduced internode length in the *ath1* mutant background (Ejaz et al., 2021), and the *pif3* and *pif4/5* mutations in *Arabidopsis* cause increased chlorophyll accumulation in leaves (Li et al., 2021). Given this, we consider the possibility that the reduced phototropic bending in the *Brachypodium elf3* mutant might be due to differences in the roles of the PIFs in *Brachypodium* as compared to *Arabidopsis.* Further work is required to determine whether this is the case.

## Materials and Methods

### Plant materials used

All *Arabidopsis thaliana* lines used in this study are of the Columbia-0 (Col-0) ecotype. The EMS-mutagenized M2 seed populations (Podolec et al., 2021) were donated by the lab of Roman Ulm (University of Geneva). The *elf3-1* (Hicks et al., 2001), *elf3-2* (Hicks et al., 2001), *lux-4* (Hazen et al., 2005), *elf4-101* (Khanna et al., 2003), *pif4-101* (Lorrain et al., 2008)*, pif5-3* (Fujimori et al., 2004)*, phyB-9* (Reed et al., 1993), *cry1-304* (Thum et al., 2001), and *phyA-211* (Nagatani et al., 1993), *elf3-2, pif4-101 pif5-3,* and *elf3-2 pif4-101 pif5-3* lines in the pTOC1::LUC background, *lux-4, lux-4 pif4-101,* and *lux-4 pif5-3* lines in the CAB2::LUC background (Nusinow et al., 2011) have been described previously. The wild type, *pif4-101,* and *pif5-3* lines in the CAB2::LUC background were generated by crossing. The *Brachypodium distachyon* accession used in this study was Bd21-3, the *elf3* mutant was described previously (Bouche et al., 2021).

### Phototropic bending assay for the mutant screen

*Arabidopsis* seeds were surface sterilized by soaking in 70% ethanol + 0.01% Triton-X-100 and agitating for 5 min, then rinsing them with 100% ethanol. Seeds were allowed to dry, and were then sown on plates containing 1/2 Murashige and Skoog (MS) medium containing 0.8% agar. The plates were then kept in the dark at 4°C for 5 days. Next, the plates were transferred to a growth chamber. The plates were mounted in black boxes that covered the sides and the bottom of the plate, so that they only received light from above (Figure S1A). The seedlings were grown in 120 µmol m^2^ s^−1^ cool white light, at 21°C, in long days (16L/8D), for 4 days. On the beginning of the 5^th^ day, the black box was rotated such that the seedlings no longer received light from above, but rather from the side (Figure S1A). Phototropic bending was assessed after 6 hours of lateral light exposure. In these conditions, wild-type seedlings do not bend towards the light-exposed side of the plate, but *phyB-9* seedlings do bend (Figure S1B). The light conditions used to grow our seedlings are further detailed in Figure S4.

### Sequencing of the *PHYB, CRY1,* and *ELF3* loci

Fragments of the *PHYB, CRY1*, and *ELF3* loci were amplified from genomic DNA using the primers listed in Table S1. These same primers were then used for Sanger sequencing of the PCR fragments. Samples were sent to Microsynth AG (https://www.microsynth.com/home-ch.html) for sequencing.

### Measuring hypocotyl length in monochromatic light conditions

Seeds were sterilized as described above, then plated on 1/2 MS medium and stratified in the dark for 5 days at 4°C. Seeds were then exposed to 4 h of white light to induce germination, before being returned to the dark for 20 h. Then, the plates were mounted vertically and either kept in the dark or exposed to different monochromatic light conditions: red (14.13 µmol m^−2^ s^−1^), far-red (4.76 µmol m^−2^ s^−1^), or blue (8.04 µmol m^−2^ s^−1^), as measured with an IL1400A photometer. Hypocotyl length was measured on the 5^th^ day after transfer to the monochromatic light conditions.

### Leaf movement assay

Seeds were sterilized, then stratified on moist filter paper in the dark at 4°C for 5 days. Then, the seeds were transferred to pots of soil. This consisted of potting soil mixed in a 2:1 ratio with vermiculite. A 0.5 cm thick layer of topsoil (Compo Sana Terreau pour semis et plantes aromatiques, from OBI) was added on top, and the seeds were placed on the surface of the topsoil. Next, a thin layer of powdered charcoal was spread over the remaining surface of the pot. A template was used to prevent the seeds from being covered by the charcoal. Then, plants were grown in white light (∼50 µmol m^−2^ s^−1^ PAR, measured with the IL1400A photometer), in long days (16 h light / 8 h dark), with a day temperature of 21.5°C and a night temperature of 20°C. On the 14^th^ day of growth, the pots were transferred to a ScanAlyzer HTS machine to for leaf tracking (for more details about the device, see Dornbusch et al., 2012). Within the ScanAlyzer, plants continued to grow in the same day/night light cycles for one day. From dawn on the next day, seedlings were kept in constant white light. Leaf movements were tracked by the ScanAlyzer machine over the course of the following 3 days.

### Circadian phototropism experiment

Seeds were sterilized and plated on 1/2 MS medium (containing 1.6% agar), which was covered in nylon mesh (Merck reference NY6H00010). The mesh was intended to reduce friction between the seedlings and the surface of the plate. The seeds were stratified at 4°C for 3 days. Then, the plates were transferred to a growth chamber. The plates were mounted vertically in black boxes such that light could only enter from the top. Seedlings were grown in combined low red:far-red (LRFR) and low blue light (LB), using cool white growth lights supplemented with far-red LEDs, and by covering the plates with a yellow filter. The light had a PAR (400-700 nm) intensity of ∼69 µmol m^−2^ s^−1^, as measured with a PG200N spectrometer. Seedlings were grown in these conditions for 4 long days (16 h light / 8 h dark). More information about the light conditions used is shown in Figure S4. On the 5^th^ day, the side of the box was opened and seedlings were exposed to lateral light, which consisted of the ambient light in the incubator, supplemented with additional blue LEDs. This gave a total lateral blue intensity (400-500 nm) of ∼25 µmol m^−2^ s^−1^. Different batches of seedlings were exposed to lateral blue light starting at different timepoints in 4-hour intervals over the course of 2 days. Phototropic bending was assessed after 4 hours of exposure to lateral blue light.

### Bending assay for *Brachypodium dystachion* plantlets

Prior to sterilizing seeds, the awns were removed to allow seedlings to grow more freely. Seeds were then surface-sterilized. First, the seeds were rinsed with water. Then, they were soaked in 70% ethanol and 0.01% Triton X-100 for 30 s, then rinsed again with water, then soaked in 1.4% sodium hypochlorite for 4 min. Finally, seeds were washed with water 3 times and placed onto moistened filter paper in petri dishes. The petri dishes were then wrapped in aluminum foil and left to stratify in the dark at 4°C for 6 days. After stratifying, seeds were transferred to 1/2 MS medium (containing 0.8% agar), with the embryo submerged in the medium and the top of the seed protruding. The seeds were exposed to monochromatic red light (∼15 µmol m^2^ s^−1^ as measured with an IL1400A photometer) for 1 hour to induce germination, and were then allowed to grow in the dark at 18°C for 4 days. To assess bending in dark-grown seedlings, the seedlings were exposed to 2 µmol m^2^ s^−1^ lateral blue light and monitored over the course of a day. To assess bending in light-grown seedlings, seedlings were first transferred to white light for 1 long day (16 h light / 8 h dark) in a black box that only allowed light exposure from above. The black box was then rotated so that seedlings were only exposed to lateral white light from the start of the second day. Seedlings were then monitored over the course of several days.

To assess which tissues contribute to bending in *Brachypodium*, we imaged seedlings used a Leica M205 FCA stereomicroscope.

### Western blots

Seeds were plated on 1/2 MS medium and stratified in the dark at 4°C. Germination was induced by exposing seedlings to white light at 21°C for 6 h, before transferring them back to the dark to grow for 4 days at 21°C. On the 5^th^ day, seedlings were either kept in the dark or exposed to 6 h of white light prior to harvesting. Seedlings were harvested into 1.5 mL Eppendorf tubes. (Dark-grown seedlings were harvested under a green light.) Then, 2x FSB buffer (20% glycerol, 4% SDS, 0.02% Bromophenol Blue, 0.125 M Tris, and 10% beta-mercaptoethanol) was added to the samples. The samples were ground manually in the buffer using small pestles, then vortexed, then heated at 95°C for 10 min. Finally, the samples were centrifuged at >16000 rcf for 10 min, and the supernatant was transferred to a fresh set of tubes.

Proteins were separated on 4-15% precast Mini-PROTEAN TGX gels (BioRad). After electrophoresis, proteins were transferred to nitrocellulose membranes using the BioRad TransBlot Turbo transfer system (mixed molecular weight program, 7 min). Total protein content in each lane was assessed by staining the membranes with Ponceau S. Next, the membranes were blocked with 1x PBS buffer containing 5% milk and 0.1% Tween-20. Blotting for PHYB was done using the BA2 primary antibody, and PHYA was done using the AA1-3 primary antibody (Shinomura et al., 1996). Both were then detected using an HRP-coupled anti-mouse secondary antibody (Promega W402B). CRY1 was probed using a primary antibody (Lin et al., 1996), and detected using an HRP-coupled anti-rabbit secondary antibody (Promega W401B). HRP-derived chemiluminescence was detected using an ImageQuant LAS4000 CCD camera system. In order to probe the same membrane with multiple antibodies, membranes were stripped between antibody treatments with a solution containing 1.5% glycine and 0.05% Tween-20, adjusted to a pH of 2.5.

### Summary of statistical methods used

Welch’s t-test was used to compare hypocotyl length and bending angle values between two populations. *p-values* from this test were calculated using the T.TEST function in Microsoft Excel (version 16.78.3 for Mac). Tukey’s HSD test was used to compare bending angles between multiple populations (See Figure 3). An alpha value of 0.05 was defined as the threshold for significance. This test was performed using the Agricolae package (version 1.3-7) in R (version 4.4.2).

## Gene accession numbers used in this study

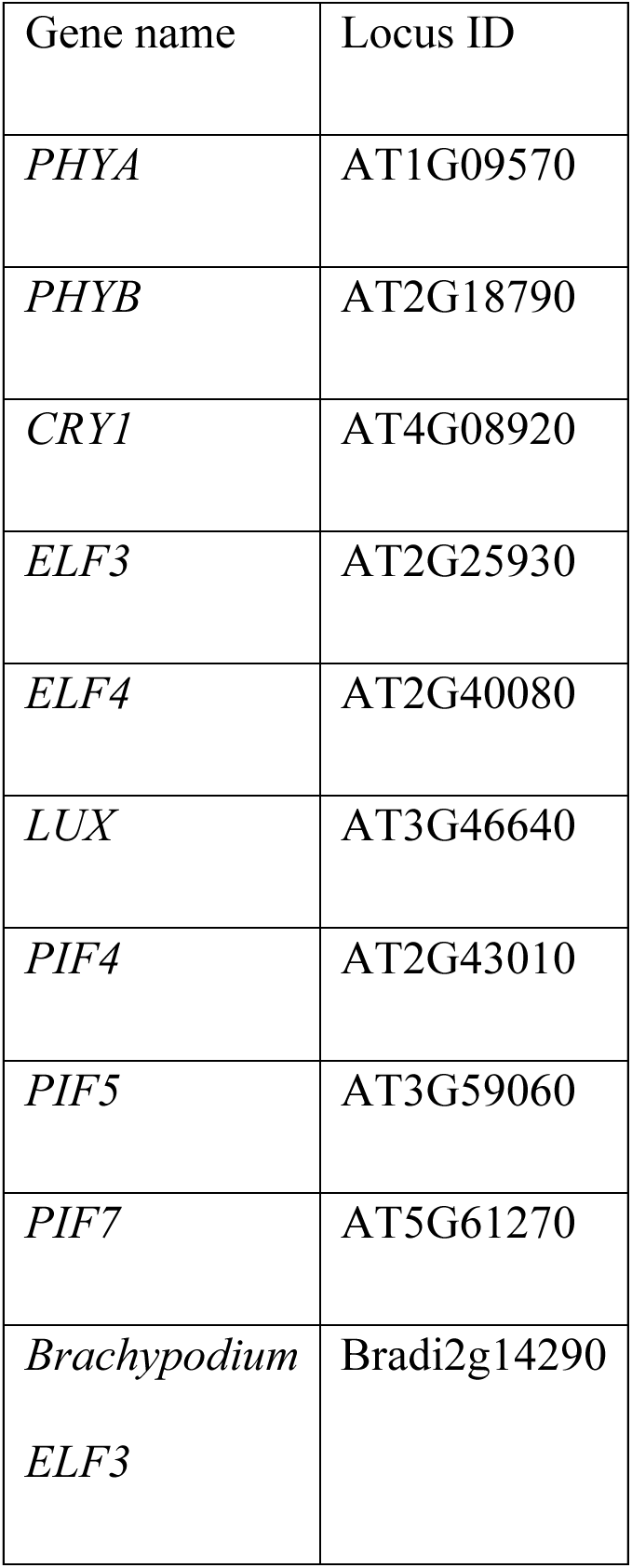

## Acknowledgement

Dmitry Nussinov (Danforth Plant Science Center) for providing seeds. We thank Roman Ulm (University of Geneva) for donating EMS mutagenized Col-0 seeds. We thank the lab of Claire Périlleux (University of Liège) for sending us the *Brachypodium* seeds. Akira Nagatani (University of Kyoto) for donating antibodies against phyA and phyB. Chentao Lin (Fujian Agriculture and Forestry University) for donating antibodies against cry1. We thank Thomas Oriol and Noémie Freymond for their technical help.

## Author’s contribution

**Conceptualization:** CF. **Formal analysis:** GC, JK, GMN, AB, CF **Funding acquisition:** CF, JK, GMN, AB. **Investigation:** GC, JK, GMN, AB **Project administration:** CF. **Validation:** GC, JK, GMN, AB, CF. **Visualization:** GC, JK **Supervision:** CF **Writing – Original Draft Preparation:** GC, CF. **Writing – Review & Editing:** GC, JK, GMN, AB, CF

## Supplemental Material

All supplemental material may be found in attached “Supplementary material” file. It comprises 1 table and 5 figures

**Figure S1.** Overview of the phototropic bending mutant screen.

**Figure S2.** Hypocotyl lengths of B111018 seedlings grown in monochromatic light conditions.

**Figure S3.** Bending in B111018 backcrosses, and genotyping B111018.

**Figure S4.** Example spectra of light conditions used in our experiments.

**Figure S5.** Western blots for PHYA, PHYB, and CRY1 in our candidate lines.

**Table S1.** Primers used in this study.

