## Supplementary figures for "Early Flowering 3 (ELF3) inhibits hypocotyl phototropism in light-grown Arabidopsis seedlings"

### ELF3 regulates positive phototropism in light-grown *Arabidopsis* seedlings.

Fankhauser.

#### Supplementary material

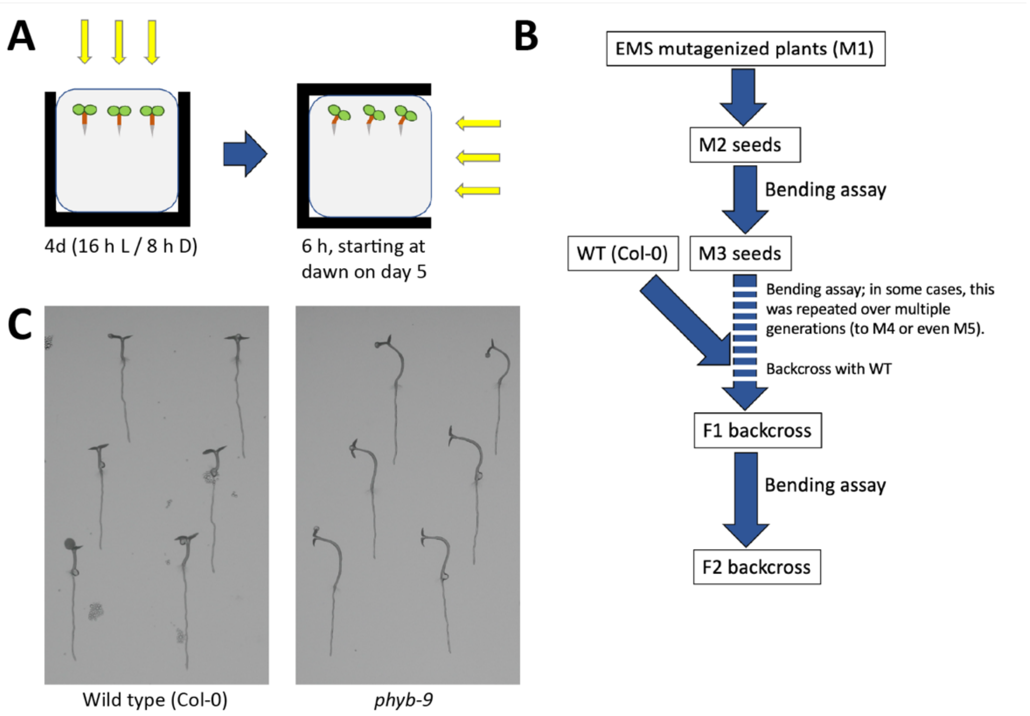


**Figure S1. Overview of the phototropic bending mutant screen. (A)** A schematic showing the setup of the bending assay in light-grown seedlings. Left: Seedlings were grown on vertically mounted plates for 4 long days (16 h light / 8 h dark) with only the top of the plate exposed to light. Right: For the bending assay, the box was rotated so that the plates were exposed to light from one side only. **(B)** A summarized workflow of the mutant screen. **(C)** Example photographs of controls (wild type and *phyB-9*) after the bending assay. *phyB* mutants bend towards the light-exposed side of the plate (to the left in this example), while the wild type does not bend.

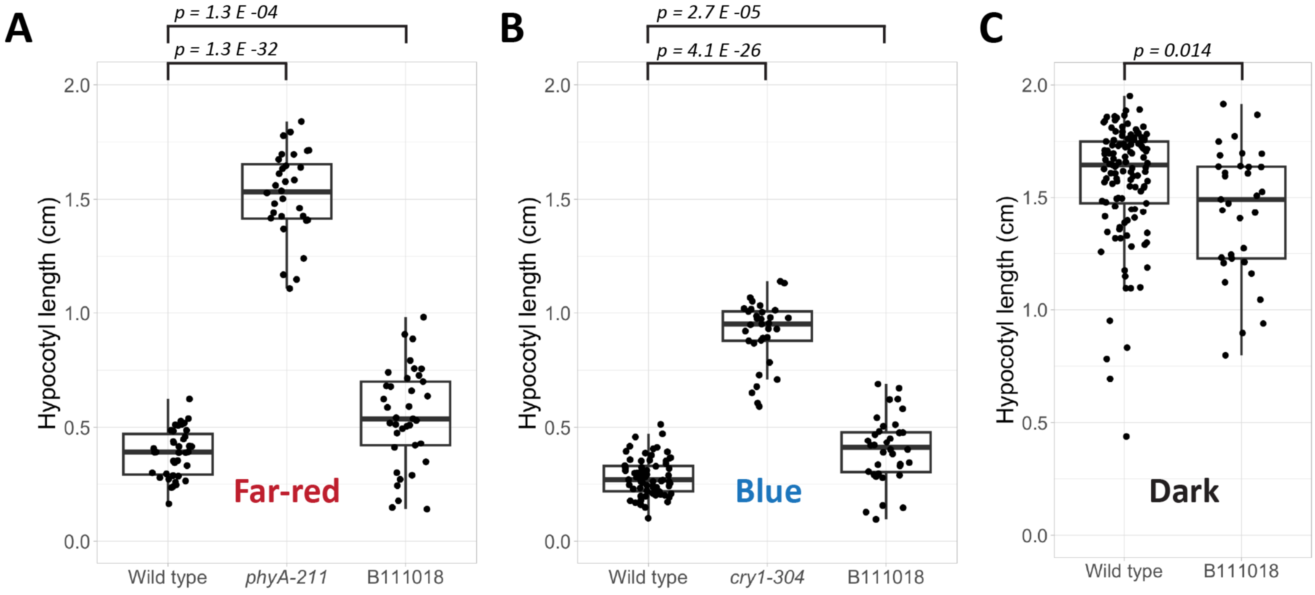


**Figure S2. Hypocotyl lengths of B111018 seedlings grown in monochromatic light conditions.** Points represent the lengths of individual seedlings; boxplots show the median length and quartiles. (A) Hypocotyl lengths in far-red light (4.76 µmol $m^2$ $s^{-1}$). (B) Hypocotyl lengths in blue light (8.04 µmol $m^2$ $s^{-1}$). (C) Hypocotyl lengths in the dark. The reported p-values are derived from Welch's t-test.

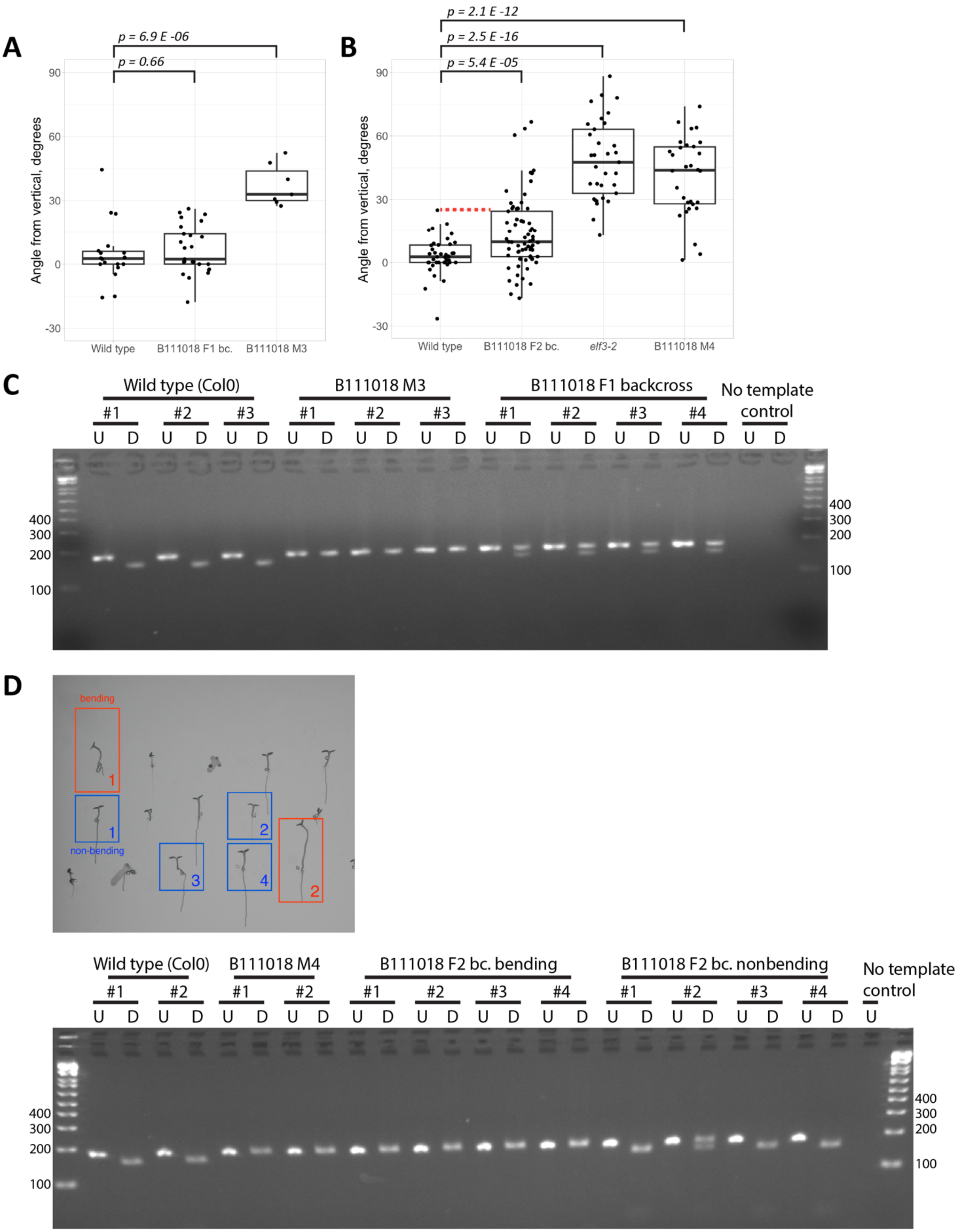
A
p = 6.9 E -06
p = 0.66
Angle from vertical, degrees
90
60
30
0
-30
Wild type
B111018 F1 bc.
B111018 M3
B
p = 2.1 E -12
p = 2.5 E -16
p = 5.4 E -05
Angle from vertical, degrees
90
60
30
0
-30
Wild type
B111018 F2 bc.
elf3-2
B111018 M4
C
Wild type (Col0)
B111018 M3
B111018 F1 backcross
No template control
#1 #2 #3 #1 #2 #3 #1 #2 #3 #4
U D U D U D U D U D U D U D U D U D U D U D
400
300
200
100
D
bending
1
1
non-bending
2
3
4
2
Wild type (Col0)
B111018 M4
B111018 F2 bc. bending
B111018 F2 bc. nonbending
No template control
#1 #2 #1 #2 #1 #2 #3 #4 #1 #2 #3 #4
U D U D U D U D U D U D U D U D U D U D U D U D U
400
300
200
100

**Figure S3 (see previous page). Bending in B111018 backcrosses, and genotyping B111018. (A)** Quantification of phototropic bending in seedlings of B111018 backcrossed to wild type (F1 generation). Points represent the bending angles of individual seedlings; boxplots show the median bending angle and quartiles. Seedlings were grown in our screen conditions for 5 days, and the bending assay began at dawn on the $6^{th}$ day. B111018 seedlings (M3 generation) are included as a reference. **(B)** Phototropic bending in seedlings of B111018 backcrossed to WT (F2 generation). *elf3-2* and B111018 seedlings (M4 generation) are included for reference. Note: p-values in (A) and (B) are derived from Welch's t-test. **(C)** Genotyping of B111018 (M3 and F1 backcross). PCR was performed with primers GC146 and GC147, using genomic DNA as a template. The PCR product was either undigested ("U") or digested with XmnI ("D"). The samples were run on a 3% agarose gel at 80V for ~30 min. The PCR product from WT plants digests to 155 + 26 bp fragments, while the 181 bp PCR product from the B111018 mutant is not digested. **(D)** Genotyping of B111018 backcross F2 seedlings. Four F2 seedlings with the increased bending phenotype and four F2 seedlings without this phenotype were selected and genotyped. Top: Sample image of a plate after the bending assay, with some of the seedlings selected for genotyping. Bending (red) and non-bending (blue) seedlings are highlighted. Bottom: Genotyping result, including the seedlings shown in the image of the plate.

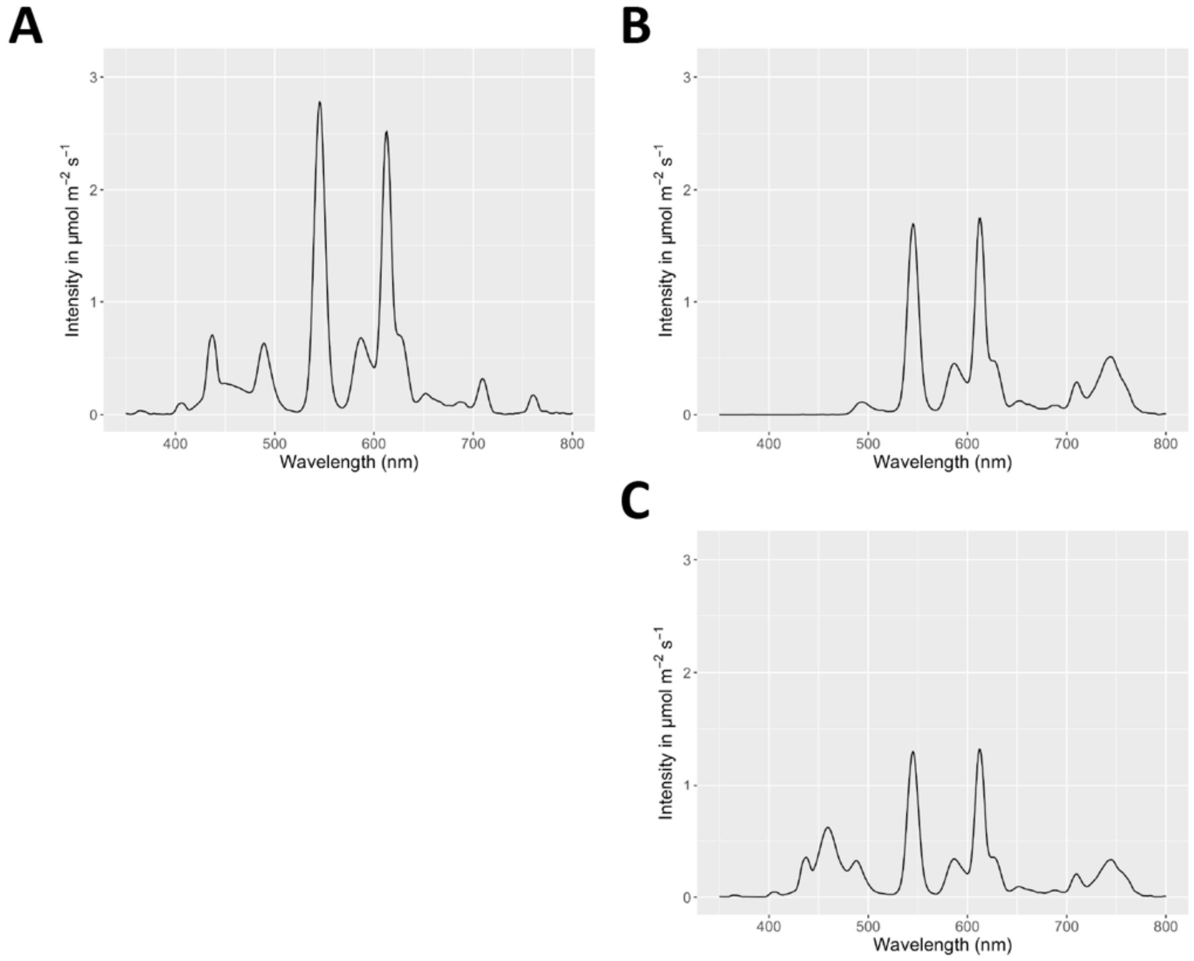


**Figure S4. Example spectra of light conditions used in our experiments. (A)** Light spectrum used for the standard bending assay. **(B)** Light spectra used for growing seedlings (top) and for inducing phototropic bending (bottom) during the circadian phototropism experiment shown in Figure 4.

**A**

| Line ID number | Blots in which present | Western blot result (M4 seedlings) |
|---|---|---|
| I04-0123 | #1<br>#4 | PHYA expressed in light and dark |
| B12-1320 | #1<br>#3 | CRY1 is not expressed |
| C09-1602 | #1<br>#4<br>#5 | PHYA expressed in light and dark |
| C12-0501 | #3 | CRY1 is not expressed |
| C06-1423 | #1<br>#2<br>#3 | Expresses CRY1 |
| C04-1719 | #1 | Expresses PHYB |
| B11-1018 | #1<br>#4 | Expresses CRY1, PHYA, and PHYB |

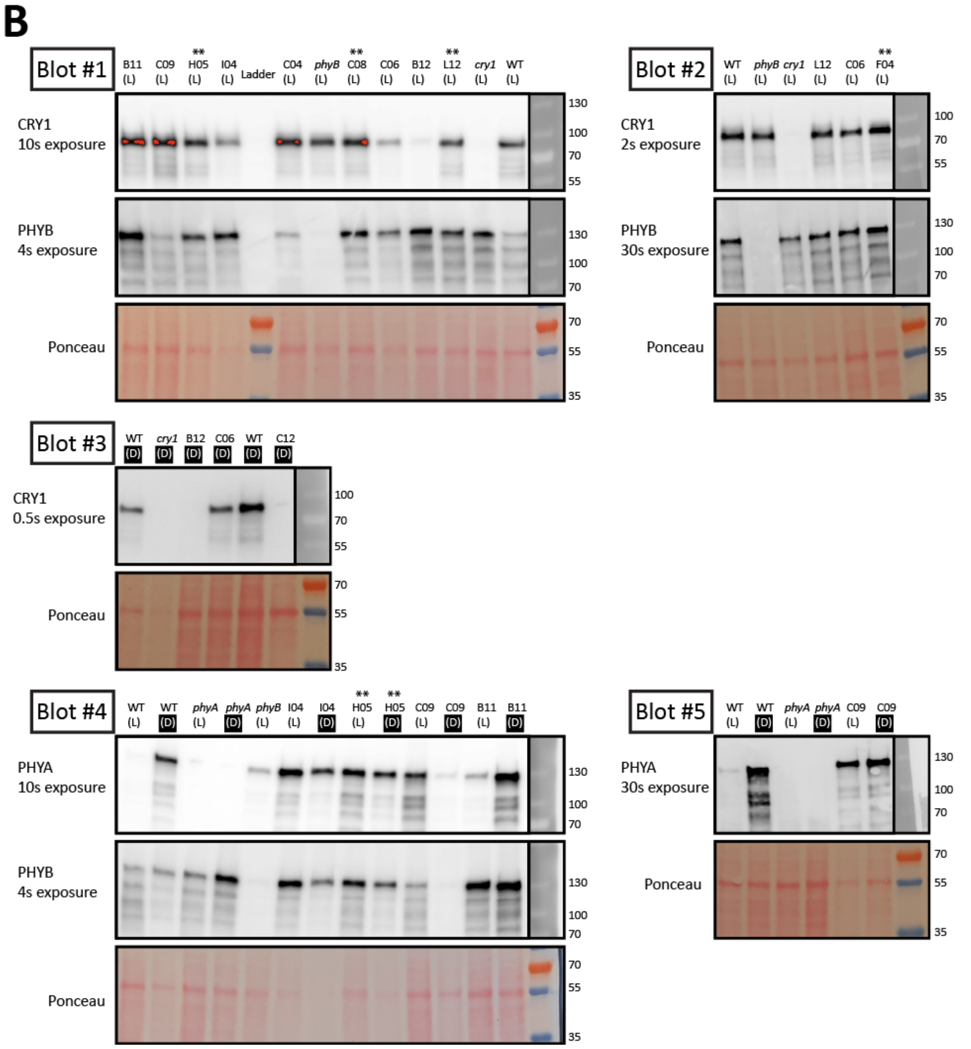


**Figure S5. Western blots for PHYA, PHYB, and CRY1 in our candidate lines. (A)** Table summarizing the candidate lines, the blots in which they are represented, and our conclusions from the western blots shown here. **(B)** The western blots. Seedlings were grown in the dark for 5 days. They were then either kept in the dark (labelled "D") or were exposed to 6h WL (labelled "L") prior to protein extraction. A color photograph of the membranes stained with Ponceau is shown to give an indication of loading. In cases where a single membrane was probed using multiple antibodies (e.g. blots 1, 2, and 4), it was stripped between probing with the first and second antibody. For more information, see Materials and Methods. Lanes marked with a "**" indicate mutant lines which were not included in our final candidate list. Orange areas in Blot #1 are indicators of overexposure marked by the ImageQuant LAS4000 imaging machine.

**Table S1:** Primers used in this study.

Part 1: Primers used for genotyping:

| Genotyping primers for: | Primers: | Expected amplicon size: |
|---|---|---|
| B111018 (*Arabidopsis elf3* allele identified in our screen) | GC146: 5'-ATGAACCAGTTTGGACATCCTGAAAATCTT-3'<br>GC147: 5'-TGCTACCAGAGATTCCCTGTG-3' | Digest the PCR product with XmnI.<br><br>WT gives bands at 155 + 26 bp.<br><br>Mutant gives a band at 181 bp. |
| *elf3-1* | OM115: 5'-TGTTGGTCAGTCTTCTCCGA-3'<br>OM116: 5'-TCCCTACTGTCATTCAAGGG -3' | Digest the PCR product with HincII.<br><br>WT gives bands at 500 + 300 bp.<br><br>Mutant gives a band at 800 bp. |
| *elf3-2* | GC41: 5'-TGGTTATTTATTCTCCGCTCTTTC-3'<br>GC42: 5'-TTGTTCCATTAGCTGTTCAACCTA-3'<br>GC43: 5'-TGAGTATTTGTTTCTTCTCGAGC-3'<br>GC44: 5'-CATATGGAGGGAAGTAGCCATTAC-3'<br><br>Note: These primers were previously described by Huang et al., 2016. | GC41 + GC42 gives a band of ~400 bp in *elf3-2,* no band in WT.<br><br>GC43 + GC44 gives 579 bp band in the WT, no band in mutant. |
| *elf4-101* | GC45: 5'-GCTTCCTATTATATCTTCCCAAATTACCAATACA-3'<br>GC46: 5'-TTCTTGAATCAAAGCAACGTTCTTC-3'<br>GC47: 5'-TTGTAGCTCTTGTCTTGCATAACAT-3'<br><br>GC45 is the standard LB2 primer used for genotyping SAIL lines (See: https://www.arabidopsis.org/abrc/pCSA110.pdf). | GC46 + GC47 gives 804 bp band in WT, no band in mutant.<br><br>GC45 + GC47 gives 563 bp band in the mutant, no band in WT. |
| *lux-4* | GC112: 5'-AACACTTAAACGACCGCCATTAGTG-3'<br>GC113: 5'-GTTCTCACGAGTTAATCCTTCAACGTTC-3' | Amplify with GC112 + GC113; digest the PCR product with BstXI.<br><br>WT gives bands at 118 + 24 bp.<br><br>Mutant gives a band at 142 bp. |
| Luciferase reporters CAB2:LUC and pTOC1:LUC | OM148: 5'-CTCACTGAGACTACATCAGC-3'<br>OM149: 5'-TCCAGATCCACAACCTTCGC-3' | A band of 107 bp occurs if a luciferase reporter is present. |
| *pif4-101* | SL42: 5'-CTCGATTTCCGGTTATGG-3'<br>SL43: 5'-CAGACGGTTGATCATCTG-3'<br>oVCG-61: 5'-TAGCATCTGAATTTCATAACCAATCTCGATACAC-3'<br><br>oVCG-61 is the standard LB3 primer used for genotyping SAIL lines (See: https://www.arabidopsis.org/abrc/pCSA110.pdf). | SL43 + oVCG061 gives 870 bp band in the mutant, no band in WT.<br><br>SL42 + SL43 gives 850 bp band in the WT, no band in mutant. |
| *pif5-3* | SL46: 5'-TCGCTCACTCGCTTACTTAC-3'<br>SL47: 5'-TCTCTACGAGCTTGGCTTTG-3'<br>oVCG-56: 5'-ATTTTGCCGATTTCGGAAC-3'<br><br>oVCG-56 is the standard LB1.3 primer used for genotyping SALK lines (See: http://signal.salk.edu/tdnaprimers.2.html) | SL46 + oVCG056 gives 860 bp band in the mutant, no band in WT.<br><br>SL46 + SL47 gives 860 bp band in the WT, no band in mutant. |

| | | |
|---|---|---|
| *elf3 (Brachypodium distachyon)* | GC154: 5'-GGTGGTTTCAGCTTCTGCAGTTGAC-3'<br>GC155: 5'-GCTGTCCCTAGATGCGTGGAATC-3' | Amplify with GC154 + GC155; Digest the PCR product with DdeI.<br><br>WT gives a band at 151 bp.<br><br>Mutant gives bands at 126 bp and 25 bp.<br><br>This *Brachypodium elf3* allele and an alternative dCAPS genotyping strategy were first described by Bouché et al., 2022. |

Part 2: Primers used for sequencing:

| **Fragment** | **Primers:** | **Expected amplicon size:** |
|---|---|---|
| *ELF3* #1 | GMN029: 5'-AGAGTCCACGTCGTCACGC-3'<br>GMN030: 5'-CACAAGGCTGCATTGAAGAGG-3' | 1103 bp |
| *ELF3* #2 | GMN031: 5'-AGGATTGAATTCTCCTGTATATATG-3'<br>GMN032: 5'-CTAGTGAACTTACTTGGCAAT-3' | 1012 bp |
| *ELF3* #3 | GMN033: 5'-AGACATTGATAATGATCGTGAA-3'<br>GMN034: 5'-ATGAGATGTCAGATATTTAACAAG-3' | 1118 bp |
| *ELF3* #4 | GMN035: 5'-TGAGTCTTATGACGATGCAG-3'<br>GMN036: 5'-TTACCAGGAGGTGGGAAT-3' | 1122 bp |
| *ELF3* #5 | GMN037: 5'-GCTCCAAATGGATATTGCT-3'<br>GMN038: 5'-GATTTCAAGGCTTTAAGACAA-3' | 881 bp |
| *CRY1 #1* | CF588: 5'-GACCAAAGGGTTTCGATTTC-3'<br>CF430: 5'-GCGGTTCTTAGAGTATTCAAGC-3' | 1067 bp |
| *CRY1 #2* | CF306: 5'-CCTTATGACCCTGAGTCTC-3'<br>CF401: 5'-CCTTGGGAACTCTATAGGAGC-3' | 1154 bp |
| *CRY1 #3* | GC29: 5'-GAGTCCGTTCTTCAAGCTGC-3'<br>GC30: 5'-GCCAACAAGTATGAGACCTGG-3' | 1283 bp |
| *PHYB #1* | GC35: 5'-TGGGCCCAATCAATCTCTCC-3'<br>GC36: 5'-ACCAGTCAAGTCCCTCACAC-3' | 1231 bp |
| *PHYB #2* | GC33: 5'-TTCTACTCGAGCGTGCTTTC-3'<br>GC34: 5'-ATTTCCGCAGTTTCCCATGG-3' | 1294 bp |
| *PHYB #3* | GC31: 5'-GATAGTTTAGGCGATGCGGG-3'<br>GC32: 5'-CCCAATCACTTCACTGCGAG-3' | 1024 bp |
| *PHYB #4* | PB16: 5'-GAATGCTTGTTCCAGCAAGG-3'<br>PB32: 5'-TGAATTCTGTGCGGATGGCG-3' | 1109 bp |
| *PHYB #5* | PB23: 5'-ACGCGATTGTAAGTCAAGCG-3'<br>AB48: 5'-TTGGCCTTTACCTCTTGATTGCGT-3' | 1259 bp |
